# GraphSeq: Accelerating String Graph Construction for De Novo Assembly on Spark

**DOI:** 10.1101/321729

**Authors:** Chung-Tsai Su, Ming-Tai Chang, Yun-Chian Cheng, Yun-Lung Li, Yao-Ting Wang

**Affiliations:** Atgenomix Taipei, Taiwan

## Abstract

**Summary:** De novo genome assembly is an important application on both uncharacterized genome assembly and variant identification in a reference-unbiased way. In comparison with de Brujin graph, string graph is a lossless data representation for de novo assembly. However, string graph construction is computational intensive. We propose GraphSeq to accelerate string graph construction by leveraging the distributed computing framework.

**Availability and Implementation:** GraphSeq is implemented with Scala on Spark and freely available at https://www.atgenomix.com/blog/graphseq.

**Supplementary information:** Supplementary data are available at *Bioinformatics* online.

## 1. INTRODUCTION

De novo assembly is one of the most fundamental problems in Bioinformatics and string graph is the powerful data representation used by many assemblers, such as SGA (Simpson and Durbin, 2011), LSG (Bonizzoni et al. 2016) and FSG (Bonizzoni et al. 2017). Comparing to de Brujin graph, string graph is more computational intensive (Bonizzoni et al. 2017). Therefore, any solution to accelerate string graph construction is desired. Apache Spark (Gupta et al. 2003) is a fast and general engine for large-scale data processing based on the MapReduce model, an in-memory computing framework for efficient data processing with parallel and distributed algorithms. In this article, we explore a novel approach, called GraphSeq, to construct string graph in parallel, based on distributed suffix array on Spark. GraphSeq can achieve 13X speedup over SGA.

## 2. METHODS

GraphSeq leverages Spark in-memory computing framework to construct string graph for de novo assembly. To fully utilize the power of distributed framework, the preprocessing step to split a big compressed file into several small ones is required. Just like Fig. 1(a), FASTQ Splitter is a python script to split a GZIP file into hundreds of small SNAPPY files. To load all of reads efficiently, GraphSeq leverages ADAM (Nothaft et al. 2015) to transform all of reads with FASTQ format to alignment records with Parquet format in Fig. 1(b). ADAM is an open source project that enables the use of Apache Spark to parallelize genomic data analysis across cluster/cloud computing environments. ADAM uses a set of predefined schemas to describe genomic sequences, reads, variants/genotypes, and features, and can be used with data in both legacy genomic file formats (e.g. FASTQ, BAM and VCF) and the co-lumnar Apache Parquet format. In Fig. 1(c), GraphSeq loads all of reads in parallel, generate the corresponding suffixes, group those suffixes with the same initial string into the same partition, and parallelly apply string graph construction algorithm by partitions. The algorithm support comprehensive configurations to adapt the different hardware specification (i.e. memory constrains, disk I/O throughput and network bandwidth) and achieve the performance improvement by fully utilizing all of resources. The detailed algorithm is described in the supplementary files.

**Fig. 1.**
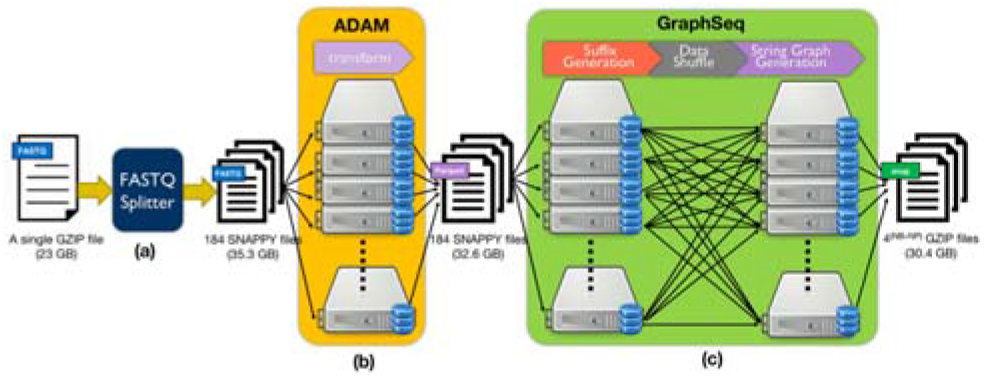
System flow. (**a**) a lightweight program to split a single big FASTQ file into several small FASTQ files. (**b**) ADAM transforms FASTQ data into alignment record in parquet format. (**c**) GraphSeq loads all of reads in parallel and generates all suffixes of the given reads. Then, each suffix will be shuffled into a specific partition by its’ prefix. After that, string graph construction can be applied within each partition.

## 3. PERFORMANCE

To demonstrate performance and scalability, the 38X WGS sample (NA12878) provided by 10X Genomics (http://s3-us-west-2.amazonaws.com/10x.files/samples/genome/2.0.0/NA12878_WGS/NA12878_WGS_fastqs.tar) is used in the paper. After applying the preprocessing steps suggested by SGA, there are 275,671,436 reads left as the benchmarking dataset and the detailed procedures is described in the supplementary files. By leveraging Spark in-memory distributed computing framework, GraphSeq has great ability of scalability and acceleration. In Fig. 2, the execution time is negatively correlated to the number of input reads when number of cores is fewer than 64. It means that the execution time will reduce by half when doubling number of CPU cores. When the number of cores is 128, it doesn’t show the same reduction ratio on execution time. There are two major reasons for this phenomenon. First, there are data skew problems on number of input suffixes and output edges when all of suffixes are partitioned by their prefix. For example, the largest (AATGGAAA) and the smallest (ACTCGACG) partitions has 42,061,542 and 3,712 suffixes, respectively. Secondly, there are 16 cores writing edge data in the single SSD simultaneously when using 128 cores and the bottleneck of disk I/O is encountered. Notice that each node of the cluster has only one SSD on Google Cloud Platform (GCP). We believe that GraphSeq maintains the same scalability on 128 or even more cores when more SSDs is added into each computing node.

**Fig. 2.**
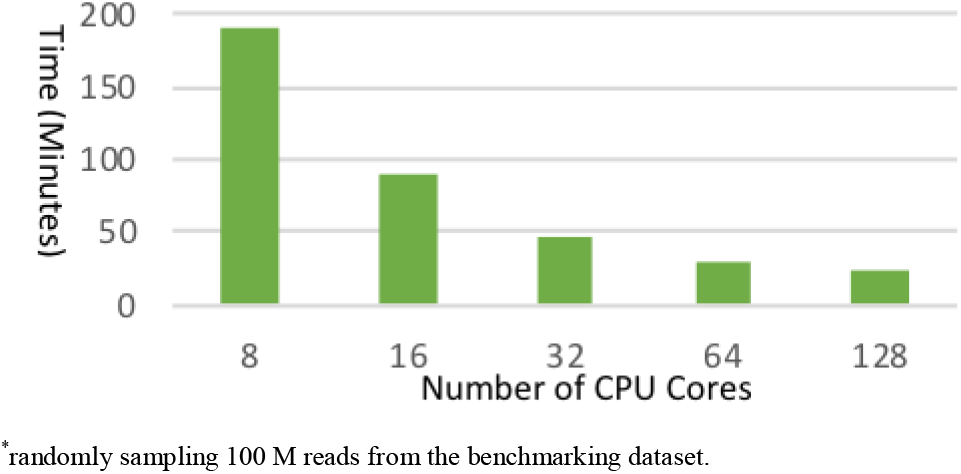
Scalability on computing resource. The cluster with 8 16-vcores computing nodes is launched on GCP.

In Fig. 3, the next experiment to demonstrate the scalability of GraphSeq is to observe the correlation between the volume of input data and the execution time. In 10M reads, the majority of execution time is the overhead of resource management from GraphSeq since 75% of tasks are executed in less than 0.6 seconds. When adding more reads to construct string graph, the execution time is increased and the linear correlation is illustrated. In the meanwhile, more edges are identified and its’ increasing rate is slightly higher than linear (214,403,795 and 627,682,220 irreducible edges in 80M and 160M reads, respectively). Therefore, the execution time is highly correlated with not only the number of input data but also the number of irreducible edges for output.

**Fig. 3.**
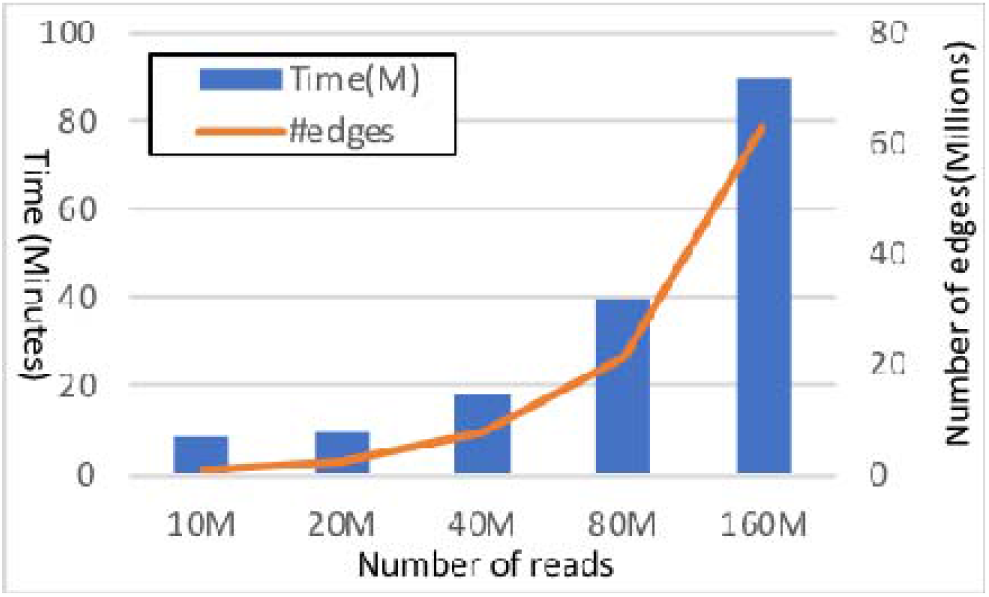
Performance evaluation on data size. The cluster with 8 16-vcores computing nodes is launched on GCP.

The performance of SGA and GraphSeq is listed in Table 1. We could not compare GraphSeq with FSG, since FSG fails to output the complete string graph with SGA although FSG performs faster than SGA. In addition, LSG is not included in the experiment since LSG is slower than SGA. Since GraphSeq leverages Spark to distribute data into multiple computing nodes in parallel for acceleration, we have to consider not only the execution time but also the total CPU hours when comparing with SGA, which is executed in a single machine. In terms of the execution time, GraphSeq outperforms SGA by 13 times faster. When the total CPU hours is considered, GraphSeq reduces around 84.6% of computational consumption in comparison with SGA. Notice that the number of edges from SGA and GraphSeq is slightly different. According to (Bonizzoni et al. 2017), SGA considers only the longer overlap when two reads have two different overlaps. GraphSeq has the same phenomenon in some situations. However, this phenomenon won’t cause any impact on the downstream process for de novo assembly or any other processes.

**Table 1.** Comparison of GraphSeq and SGA.

| Method | Time(Minutes) | Total CPU Hours | #edges |
| --- | --- | --- | --- |
| SGA* | 1,639 | 874.1 | 1,362,829,548 |
| GraphSeq** | 126 | 134.4 | 1,362,829,567 |

\*SGA is executed on the instance with 32-vCores on GCP.
\*\*GraphSeq is executed on the 8-nodes cluster on GCP. Each node has 16 vCores and 60 GB of Memory.

## 4. CONCLUSIONS AND FUTURE WORK

We present GraphSeq, a tool implementing on Apache Spark, to construct string graph in parallel and achieve 13x speedup over SGA. In future, we will provide the corresponding tools on Spark to deliver a streamline pipeline for de novo assembly, including error correction and contig assembly.

## Acknowledgements

We thank Jen-Ming Chung for early development.

